# Directed evolution of the PcaV allosteric transcription factor to generate a biosensor for aromatic aldehydes

**DOI:** 10.1101/689232

**Authors:** Leopoldo F. M. Machado, Andrew Currin, Neil Dixon

## Abstract

**Background:** Genetically encoded biosensors are useful tools for the detection of metabolites and industrially valuable molecules, and present many potential applications in biotechnology and biomedicine. However, the most common approach to develop biosensors relies on employing a limited set of naturally occurring allosteric transcript factors (aTFs). Therefore, altering the substrate specificity of aTFs towards the detection of new effectors is an important goal.

**Results:** Here, the PcaV repressor, a member of the MarR aTF family, was used to develop a biosensor for the detection of hydroxyl-substituted benzoic acids, including protocatechuic acid (PCA). The PCA biosensor was further subjected to directed evolution to alter its substrate specificity towards vanillin and other closely related aromatic aldehydes, to generate the Van2 biosensor. Substrate recognition of Van2 was explored *in vitro* using a range of biochemical and biophysical analyses, and extensive *in vivo* genetic-phenotypic analysis was performed to determine the role of each amino acid change upon biosensor performance.

**Conclusions:** This is the first study to report directed evolution of a member of the MarR aTF family, and demonstrates the plasticity of the PCA biosensor by altering its substrate specificity to generate a biosensor for aromatic aldehydes.

## Background

Biosensing is an essential mechanism widely employed in nature to perceive environment conditions; such as the presence of certain intracellular or extracellular metabolites; nutrients, metal ions and stress signals [1, 2]. Sensing is commonly associated with the regulation of gene expression at the transcriptional level mediated by an allosteric transcription factor (aTF). The aTF is a protein composed of an effector binding domain (EBD) that senses an effector molecule and a DNA binding domain (DBD) that interacts with a specific DNA operator sequence. Binding of an effector to the EBD leads to an allosterically induced conformation change affecting the interaction of the DBD with the DNA, hence modulating promoter accessibility and transcription initiation [1, 3, 4]. Generally, aTFs can be divided in repressors or activators. Repressors bind to a DNA operator blocking the RNA polymerase (RNAP) promoter binding, subsequent effector binding to the EBD leads to derepression allowing transcription initiation. Activators bind to DNA operator sequences permitting RNAP binding to the promoter to active transcription, often requiring aTF contact with the RNAP α-subunit [4, 5].

An aTF and operator sequence can be harnessed and used in the design of genetic circuits to control expression of a reporter gene, generating a genetically encoded biosensor for the cognate effector [6, 7]. The reporter is usually a fluorescent protein, an enzyme or a toxic gene allowing a range of applications including high-throughput screening methods [8], adaptive laboratory evolution [9, 10], protein directed evolution [11], metabolic pathway optimisation [12, 13] and metabolite monitoring [14]. Gene regulation tools based on aTFs have revolutionised biology. Canonical aTFs like LacI [15], AraC [16] and TetR [17] have been applied as inducible gene expression for decades. More recently, aTFs have been employed as biosensors for the detection of other molecules including industrially valuable aliphatic [12, 18, 19] and aromatic [20–22] chemicals. Although aTF biosensors have been broadly used, the repertoire of effectors with a known aTF is limited [7]. Therefore strategies to generate new biosensors for desired molecules have been pursued [23, 24].

The directed evolution of the aTF to change its substrate specificity is a versatile alternative to develop new biosensors [25–29]. Nonetheless, engineering transcription factors is inherently challenging due to their allosteric nature. Small changes in the EBD can affect the stability of the DBD and the performance of the whole system [30]. Examples of aTFs reengineered to accommodate alternative substrates include LacI [31, 32], TetR [33, 34], AraC [26, 35, 36] and TrpR [29]. In general, these systems are very well characterised biochemically, structurally, and mechanistically, which greatly facilitates the directed evolution process.

PcaV is an aTF repressor of the MarR family involved in the genetic regulation of the catabolic genes for protocatechuic acid (PCA) in *Streptomyces coelicolor* [37, 38]. This family of aTFs include examples for the sensing of environmental signals including antibiotics, toxic molecules, reactive species [39] and small molecule metabolites [40]. In contrast to most of the aTFs previously employed in directed evolution studies, relatively less is known about the allosteric mechanism of the MarR aTFs [41, 42].

Aromatic aldehydes are important molecules in an industrial context, as they can be used as chemical building blocks and have major applications in the flavors industry, especially benzaldehyde and vanillin [43, 44]. Furthermore, some aromatic aldehydes can be produced from lignin, a major component of plant biomass, therefore providing an alternative sustainable source for these key chemical building blocks away from fossil-derived sources [45]. The presence of a large market added to low yields from natural extraction and negative perception towards synthetic chemistry methods have recently increased the demand for biotechnological production of aromatic aldehydes [46, 47]. Therefore, improvements in the bioproduction and application of these molecules would be highly desirable and valuable, and would be greatly enhanced by the development of new biosensor systems to expand the biotechnological tools available for discovery and innovation in the area.

In this study a PCA biosensor, based on the PcaV aTF, was developed and a wide range of candidate effectors were assessed. A directed evolution strategy of PcaV was then employed to alter the substrate specificity of the PCA biosensor to detect a range of small aromatic aldehydes including vanillin. The resultant Van2 biosensor was functionally and biochemically characterised and further mutations were introduced to explore the role of each amino acid change upon the biosensor performance.

## Methods

### Primers, Plasmids and Strains

The DNA oligos used in this study for constructions and sequencing are presented in **Table S1**; oligos used to generate the PcaV DE libraries are described in the **Table S2**. Plasmids details are described in **Table S4**. Strains are described in **Table S5**. The DNA parts with the gene for *pcaV* and the promoters P_LV_ and P_PV_ were synthetized by GenArt. *pcaV* was cloned into a p15 plasmid flanked by a pLacI constitutive promoter and a rrnB1 terminator by restriction digestion of the *NdeI* and *XbaI* sites. P_LV_ and P_PV_ were cloned into a pET44-eGFP plasmid [22] upstream to an *eGFP* gene by restriction digestion using *NdeI* and *SphI* sites. Cloning transformation was made into *E. coli* DH5-alpha (*NEB*), transformation for induction test was made into *E. coli* BL21 (*NEB*).

### Chemicals

The chemicals used in this study with the description and suppliers are presented in **Tables S6** and **S7**. Stocks were prepared dissolving the chemicals in dimethyl sulfoxide (DMSO) at 100 mM concentration. Dilution of the stocks was made dissolving them in water, or DMSO for hydrophobic chemicals.

### Biosensor cultures growth and induction assays

*E. coli* (BL21, BW25113 or RARE [48] strains) were co-transformed with the required plasmids for the PCA biosensor system, PcaV library (Table S4) or Van2 biosensor system and grown in LB media supplemented with chloramphenicol (30 μg/mL) and ampicillin (100 μg/mL). The induction assays were performed by the addition of the tested substrates or effectors to the *E. coli* biosensor cultures freshly grown to exponential/mid-log stage (cell density ∼ OD 0.6) in deep well plates. The eGFP expression was monitored following three hours of induction at 37°C with shaking at 1000 RPM. Cells were centrifuged, washed and re-suspended with PBS buffer. The expression output was then analysed by monitoring the fluorescence normalised to cell density (RFU/OD_600_) in a plate reader (*ClarioStar*).

### PcaV Libraries design and construction

The PcaV libraries were constructed using the SpeedyGenes gene synthesis method [49, 50]. PcaV WT and library genes were designed with the GeneGenie software [51], allowing the generation of WT or variant libraries by overlap-extension PCR (OE-PCR) using 10 overlapping oligonucleotides. Terminal primers also encoded homologous cloning sequences to allow the ligation of the final PCR product into the p15 plasmid backbone. Wild type and variant (NNK codon) oligonucleotides were designed for each of the 7 selected amino acid positions, His9, His21, Trp25, Tyr38, Ile110, Met113, Asn114 (**Table S2**). Different combinations of the oligos in the **Table S2** were then used to generate libraries of PcaV mutants completely randomized for 3 amino acids (libraries PcaV DE L1; L2; L3 and L4) (**Table S3**).

PcaV WT and PcaV libraries were made in two steps. First an OE-PCR was performed employing the selected set of 10 oligonucleotides, according to the desired product. Amplification was made using a Q5 Hot-Start High-Fidelity DNA Polymerase (*NEB*) with 35 cycles of Denaturation (98°C, 20 seconds), Annealing (65 °C, 15 seconds) and Extension (72 °C, 30 seconds). Secondly, the product band was gel purified with the MinElute Gel Extraction Kit (*QIAGEN*) and an endonuclease correction step was performed with the Surveyor^®^ Mutation Detection Kit (IDT) as previously described at [49]. Finally, a second PCR step was performed to amplify the Surveyor treated template with the gene’s end primers (PcaV DE WT 1F and PcaV DE WT 10R). The final PCR product was cloned into a p15 plasmid flanked by a pLacI constitutive promoter and the rrnB1 terminator. Ligations were made by isothermal assembly with the NEBuilder HiFi DNA assembly (*NEB*) using the protocol suggested by the supplier.

Libraries were transformed into *E. coli* DH5-alpha, grown in high volume flasks until OD ∼1.0 and plasmid extracted. A fraction of the grown cultures was spread on LB agar plates for colony counting and 10 colonies from each library were sequenced for the quality control. The experimental library size ranged from 2.5 × 10^4^ to 8.0 × 10^4^ (**Table S3**), which is generally determined by ligation and transformation efficiency. The accuracy of the library construction, shown as percentage of perfect gene sequences, was > 60% for all the libraries (typically random indels). All the 10 sequences for each library had different nucleotides on the variant codon (**Table S3**).

The PcaV DE libraries were transformed into electrocompetent *E. coli* BW25113 (K strain) cells harbouring the reporter plasmid p44pPVeGFP. The libraries 1 to 4 were combined to compose the library PcaV DE L1234 and the screening step was followed.

### Fluorescence Activated Cell Sorting (FACS) counter-selection

Cultures of *E. coli BW25113* without plasmids (Negative control, C-) or transformed with plasmids containing eGFP reporter (Positive control, C+); the PCA WCB system (Biosensor control); and the PcaV DE L1234 library (Tests) were grown overnight in LB media supplemented with antibiotics. Cultures were diluted (1:100) into fresh media, grown until OD∼0.6 and induced with vanillin or PCA (where appropriate) for 3 hours.

Cells were centrifuged, washed with PBS and analysed using FACS (*Sony SH800*). Populations were sorted according to the screening method for the enrichment of mutants with altered substrate specificity. Four rounds of FACS counter-selection were made alternating negative sorting, where uninduced libraries were used to sort events on the bottom of the GFP fluorescent histogram; and positive sorting, where libraries induced with the desired effector were used to sort events on the top of the GFP fluorescent histogram.

In the 1^st^ round, negative sorting was performed, collecting the population on the bottom 5% of the histogram. At least 2.5 × 10^6^ events were passed through the flow cytometer, which correspond to ∼20-fold the experimental library size (1.3 × 10^5^), ensuring the visualization of all variants. In the 2^nd^ round, libraries were induced with the new candidate effector vanillin at 1 mM followed by positive sorting of the top 5% of the histogram. At least 10-fold the initial library size was passed through the flow cytometer. In the 3rd round, the bottom 50% was collected, again passing a number of events corresponding to 10-fold the initial library size. And finally, a 4^th^ round was performed inducing the libraries with vanillin at 1mM and sorting the top 5% of the histogram (**Figure S2**). After the last round, the sorted cells were spread on LB agar plates supplemented with ampicillin (100 μg/mL) and chloramphenicol (30 μg/mL). The same procedure was performed using the parent effector PCA in parallel as a control.

A 96-well plate based screening assay was performed with 48 colonies of each sorted library testing them with the biosensor induction assay. Cultures were induced with 1.0 mM of the corresponding inducer or water. The fluorescence was measured and the ratio induced/uninduced was used to select for responsive mutants.

### GC-MS analysis

Supernatants of cultures uninduced or induced with 250 μM, 500 μM and 1mM of vanillin and vanillic acid were collected and frozen at −80°C. Organic extraction was followed. 400 uL of chemical standards and defrosted samples were collected and the internal standard C9 fatty acid was added to final concentration of 500 μM in each one. 50 μL of 5 M HCl was added and the tubes were vortexed for 2 min. 400 μL of methyl-tert-butyl ether (MTBE) was added, the tubes were vortexed again for 2 min and centrifuged at 11,000 G for 5 minutes. Around 300 μL of the organic layer was collected and transferred to 1.5 mL tubes containing small amounts of MgSO_4_ powder. The pellet was mixed to make sure all the water was adsorbed and centrifuged at 11,000 G for 2 minutes. 180 μL of the supernatant was added to HPLC vials (*Sigma*) filtering it to make sure no trace of MgSO_4_ was present in the sample. Derivatisation was made by adding 20 μL of N,O-bis(trimethylsilyl) + 1% trimethylchlorosilane (BSTFA + 1% TMCS) and then incubating the vials at 50°C for 30 min. Analysis was followed in GC-MS (*7890B GC System /5977B MSD Agilent Technologies*).

### Sub-cloning, protein expression and purification

Sequences encoding PcaV WT and Van2 mutant were PCR amplified from the plasmids p15pcaVWT and p15van2 (**Table S4**) inserting a N-terminal TEV cleavage tag and cloned into a pET28a backbone in front of the His-tag using the NEBuilder HiFi DNA assembly method (*NEB*).

The genes were both expressed with the same conditions using *E. coli* BL21 (DE3) (*NEB*) leading to the PcaV and Van2 proteins. An overnight pre-culture of LB broth (*Formedium*) supplemented with Kanamycin 50 μg/mL was inoculated from - 80°C glycerol stocks. The expression cultures were prepared diluting the overnight cultures into LB media (1:100) and incubating at 37 °C, 180 RPM until OD 0.8. Induction was made with IPTG 0.5 mM for 20 hours at 18°C. Cells were harvested by centrifugation at 5000 RPM, 4C for 1 hour.

The pellets were re-suspended with lysis buffer (25 mM Tris-HCl, 500 mM NaCl, 10 mM imidazole, pH 7.5) supplemented with EDTA-free protease inhibitor (Roche) and 2.5 U/mL Benzonase nuclease (Sigma). Cell lysis was made using high-pressure with a cell disruptor and centrifuged at high speed (40 KG) to remove the debris. The lysate was loaded into a column of Ni-NTA Agarose beads (Qiagen) and the protein was purified using increasing concentrations of imidazole. The purified fractions were analysed on a SDS-PAGE, the cleaner fractions were mixed and dialysed.

The His-TEV tag was cleaved using TEV protease (Sigma). The reaction was made overnight at 4°C into dialysis buffer for TEV (25 mM Tris-HCl, 500 mM NaCl, 10 mM imidazole, 1 mM DTT, 0.5 mM EDTA, pH 8.0). The DTT and EDTA were removed immersing the cleaving reaction into dialysis buffer (25 mM Tris-HCl, 500 mM NaCl, 10 mM imidazole, pH 7.5) for one hour. The TEV cleaved protein was obtained by a second Ni affinity purification step to remove the His-TEV peptide and the TEV protease. Further purification step was performed by size-exclusion using a Superdex 200 10/300 GL (*GE Healthcare*) column on an AKTA plus system. This final protein was analysed by SDS-PAGE and concentrated.

### Biophysical characterisation

Freshly prepared proteins were used to analyse the biophysical properties and dimerization by size-exclusion chromatography coupled to multi-angle light scattering (SEC-MALS) analysis. The samples of PcaV and Van2 (0.75 mL at 1.5 mg/mL) were loaded in a size exclusion column Superdex 10/300 GL (*GE Healthcare*) connected to a Wyatt DAWN Heleos II EOS 18-angle laser photometer coupled to a Wyatt Optilab rEX refractive index detector. Data analysis was performed using the Astra 6 software (Wyatt technology Corp.)

Surface Plasmon Resonance (SPR) experiments were performed using the Biacore T200 system (*GE Healthcare*). The PcaV and Van2 proteins obtained from the SEC peak were diluted in SPR buffer (10 mM Hepes, 150 mM NaCl, pH 7.4). The PV Bio DNA probe with a biotin tag attached to the 5’ end was ordered from IDT (Table S1). The SPR chip GE CM5 (GE) was used with SPR buffer (10 mM Hepes, 150 mM NaCl, pH 7.4). In a constant flux, neutravidine was immobilised on the chip’s covalent dextran matrix. Blocking of the non-bound area was made with ethanolamine present in the quenching buffer. Around 100 nM of the 5’ Bio-PV probe was used to bind the neutroavidine and cover the chip. Decreasing concentrations of PcaV or Van2 were tested to determine the affinity constant to the DNA probe. Concentrations of PcaV from 2 μg/mL to 240 μg/mL (0.94 nM to 60 nM) and Van2 ranging from 20 μg/mL to 240 μg/mL (9.4 nM to 600 nM) were used.

### Electrophoretic Mobility Shift Assay (EMSA)

EMSA was performed with an infra-red based method using *LI-COR* EMSA Kit (*LI-COR*. Reactions were prepared adding stock Binding Buffer to obtain the final concentrations (50mM Tris–HCl, 50mM KCl, 5mM MgCl2, 1mM EDTA, 5% (v/v) glycerol, 0.1 ug poly (dI-dC), 1mM dithiothreitol (DTT) in the reaction tube. The DNA PV probe infra-red duplex was purchased from IDT containing one palindromic sequence from the chimeric promoter P_PV_ flanked by 6 nucleotides (32mer in total) and a 700nm fluorophore tagged on the 5’ end of each primer (IR700-5’-TTGACT<u>ATACTCAGTGCCCTGACTAT</u>GATACT-3’). The control probe Rnd Probe IR700 duplex with the same nucleotides completely randomized was also purchased from IDT. Novex TBE - DNA retardation gels (6%) – (*ThermoFisher*) were used for the electrophoresis.

Binding assays were performed adding stocks solutions of binding buffer, nuclease-free water, DNA probe (500 pM) and Van2 (500 nM) or PcaV (250 nM) proteins, according to the test order. Binding was performed by incubating the reaction tubes for 20 min at 30°C. Substrates (2.5 mM to 12.5 mM) were added to the binding reactions and incubated again for 30 min at 30°C. The final reactions were mixed with 10X Orange Loading Dye (*LI-COR*), applied to the gel and electrophoresis was performed for 45 minutes at 100 volts. Imaging was made straight away using a 700 nm excitation laser in an Odyssey imager (*LI-COR*).

### Site-directed mutagenesis

Site-directed mutagenesis was used to generate point mutation of the PcaV to Van2 aTF as well as to generate further mutants from Van2. A reverse constitutive primer (van2 PM RV – Table S1) was designed to amplify the gene downstream of the codons for amino acids 110, 113 and 114. Forward primers were designed to include one or two codon mutations keeping an overlap sequence with van2 PM RV oligo of at least 12 nucleotides (**Table S1**). Site-directed mutagenesis PCR was made with Q5 Hot-Start High-Fidelity DNA Polymerase (*NEB*) to amplify the p15van2 plasmid incorporating the mutations. The PCR product was ligated using the NEBuilder HiFi DNA assembly (*NEB*).

### Modelling and docking

Homology models of Van2, and related variants, were generated with SwissModel [52] (https://swissmodel.expasy.org/) using the crystal structure of PcaV complexed with PCA (4FHT) as the template. PcaV was generated for re-docking manually deleting PCA and water molecules, except the coordinated water (HOH13), from 4FHT. All protein structures were energy minimised using either GROMOS96 force field tool inside SwissPDB-Viewer or Amber14SB force field inside Chimera, leading to similar outputs after docking. The effectors (PCA and vanillin) were prepared in ChemDraw and energy minimised using MM2 force field in Chem3D. Docking of PcaV (re-docking), Van2, and PcaV to Van2 mutants with PCA and vanillin was made with AutoDock Vina (version 1.1.2) [53]. The docking was performed with a flexible setting, which is based on selecting both specific amino acids in the binding-pocket and the effector to rotate during the docking. The 3 amino acid positions mutated from PcaV to Van2 (110, 113 and 114) and effectors (PCA and vanillin) were allowed to rotate. Docking results listed the 10 best conformational states in a score based on the lowest ΔG_binding_ in kcal/mol.

## Results

### Construction and performance of the PCA Biosensor

A biosensor was developed for the detection of protocatechuic acid (PCA) using the PcaV allosteric transcription factor from *Streptomyces coelicolor*. PcaV interacts with its DNA operator region (Oi) regulating the pcaI gene via a repression mechanism, which is abolished upon binding by hydroxyl-substituted benzoic acids, including PCA [38]. To develop the reporter component of the biosensor we employed a chimeric promoter engineering strategy [22]. Here the PcaV-Oi sequence from *S. coelicolor* [38] was used to design two chimeric promoter-operator sequences based on phage promoter architecture, to generate a lambda phage chimeric promoter operator (P_LV_), and a phage T7A1 chimeric promoter operator (P_PV_) [54] (**Fig. S1A**). The engineered promoters were then used to generate two biosensor designs placing each one upstream of an eGFP reporter gene. Based on this design, the reporter expression should be repressed by PcaV, and then derepressed upon PCA binding to PcaV, permitting transcription, expression of the reporter protein and generation of a fluorescent signal (**Fig. 1A**).

**Figure 1:**
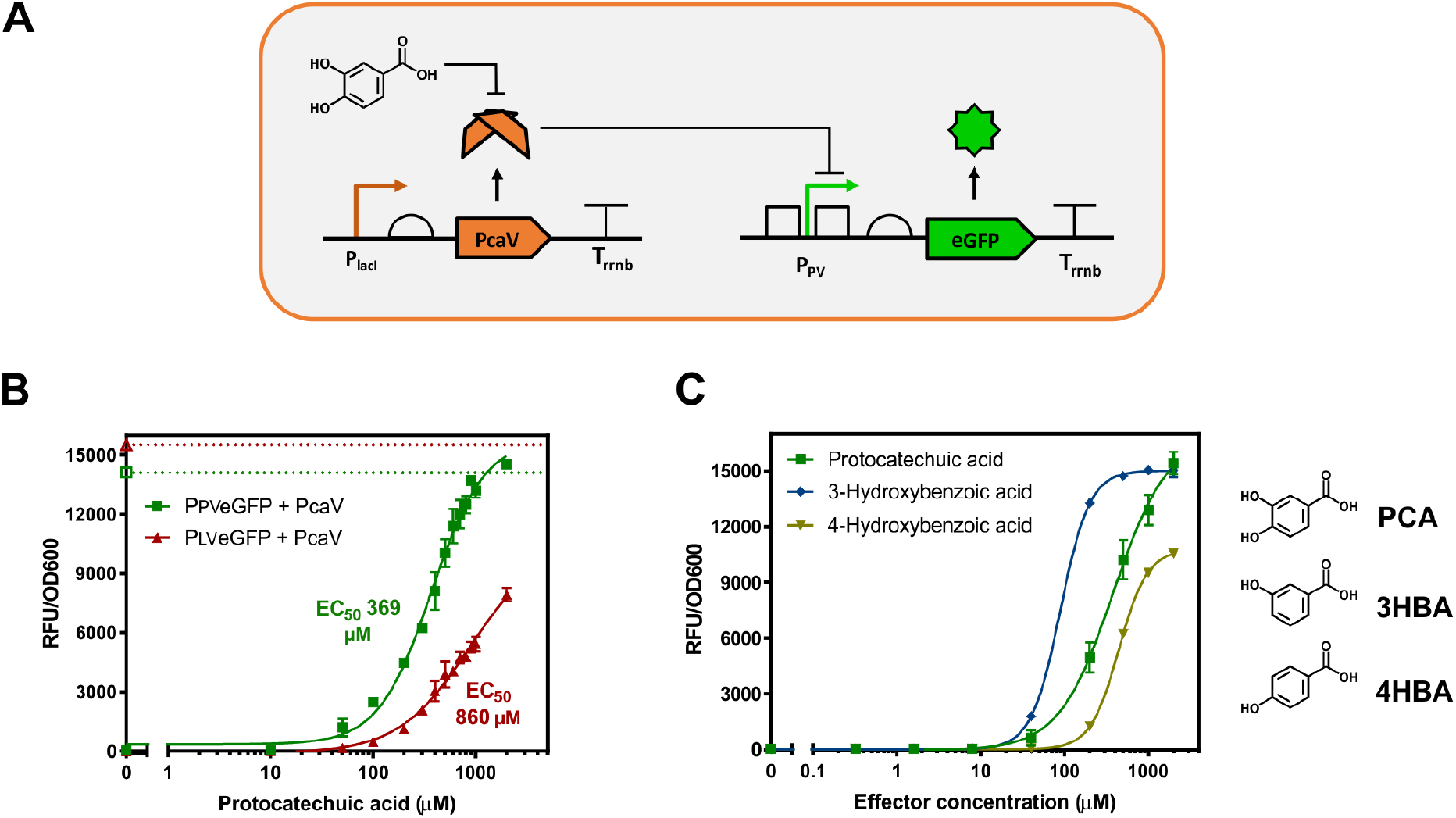
The PCA Biosensor in *E. coli*. **A**. Representation of the biosensor for detection of protocatechuic acid (PCA) is shown. Engineered chimeric promoter-operator (P_PV_) inserted upstream from the eGFP reporter gene. PcaV binds to the promoter-operator sequence(s) repressing the expression of the reporter gene. Upon PCA binding to PcaV derepression occurs generating a fluorescent signal. **B.** PCA biosensor dose response curves. eGFP gene expression data in the absence (empty shapes) and presence of the PcaV repressor (filled shapes), for the P_LV_ (red triangles) and P_PV_ (green squares) versions of the PCA biosensor in *E. coli*. **C.** Titration curves for active screened compounds using the P_PV_ version of the PCA biosensor in *E. coli* and their respective effector. The fluorescent gene expression normalized to cell density (RFU/OD_600_) is shown relative to increasing concentrations of the effector. Each value represents the mean and standard deviation of 3 biological replicates.

The plasmids permitting constitutive expression of PcaV (p15PcaV) and regulated expression of the eGFP reporter (p44P_PV_ or p44P_LV_), which encode the biosensor system, were co-transformed into *E. coli* cells and their performance was analysed with increasing concentrations of PCA (**Methods**). The P_PV_ biosensor displayed the highest signal output, dynamic range (∼150-fold) and greatest sensitivity EC_50_ of 369 μM, whilst the P_LV_ biosensor showed a reduced signal, dynamic range (∼90-fold), and sensitivity EC_50_ of 860 μM (**Fig. 1B**). The best version of the PCA biosensor, the P_PV_, was selected and used for all subsequent experiments.

Following initial performance assessment, a substrate specificity screen was conducted to identify additional effectors for the PCA biosensor. In total 29 aromatic molecules with chemical structures and functional groups, related to PCA, were selected and assessed with the PCA biosensor (**Table S6**). The initial screening was performed with fixed concentration of each candidate effector (1 mM) (**Fig. S1B**), the identified active effectors were then assessed in a full titration screen (**Fig. 1C**). Three active effectors were able to generate an increased output signal: protocatechuic acid or 3,4-dihydroxybenzoic acid (PCA), 3-hydroxybenzoic acid (3HB) and 4-hydroxybenzoic acid (4HB). These three effectors contain a very similar chemical structure, with the presence of a hydroxyl group in either the 3 or 4 position on the aromatic ring for 3HB and 4HB respectively, or both 3 and 4 for PCA. Interestingly, similar aromatic molecules did not display activity. For example, the presence of an additional hydroxyl group on the position 2 (gallic acid), presence of a methoxy group on position 3 (vanillic acid) and absence of substitutions (benzoic acid) resulted in no activity demonstrating a narrow substrate specificity pattern of the PCA biosensor (**Fig. S1B**).

The large dynamic range and narrow substrate specificity of the PCA biosensor creates sensing capabilities for the detection of hydroxyl substituted benzoic molecules, which are commonly present as intermediates in the microbial catabolic degradation pathways of lignin [55, 56]. Moreover, these performance characteristics make the PCA biosensor an ideal system to explore the potential to alter its substrate specificity using directed evolution strategies [27, 57].

### PcaV Directed Evolution

A directed evolution strategy was employed in order to assess the plasticity of the PcaV aTF and to explore whether biosensors with altered substrate specificity could be developed from the PCA biosensor.

The structural information regarding the PcaV-PCA interaction has been previously described (PDB 4G9Y, 4FHT) [38]. Residues from both dimers interact with PCA in each binding-pocket: H21(A), W25(A), S35(A), Y38(A), A39(A), I110(A), M113(A), N114(A) from monomer A and L6(B), H9(B), G11(B), H12(B), R15(B) from monomer B (**Fig. 2A**). The amino acids on the lower part of the binding-pocket are associated with the repression mechanism, especially R15 [38]. Based on this information, a structure-based directed evolution strategy was proposed. The amino acids known to interact with the 3-hydroxy and 4-hydroxy groups, on the upper side of the effector binding-pocket, were selected for mutagenesis, whereas the amino acids on the lower side were maintained (**Fig. 2A**). DNA libraries encoding PcaV were designed and assembled (PcaV DE L1; L2; L3 and L4), using the NNK codon set, at positions corresponding to the 7 amino acid residues H9, H21, W25, Y38, I110, M113, N114 (**Table S1-2**).

**Figure 2:**
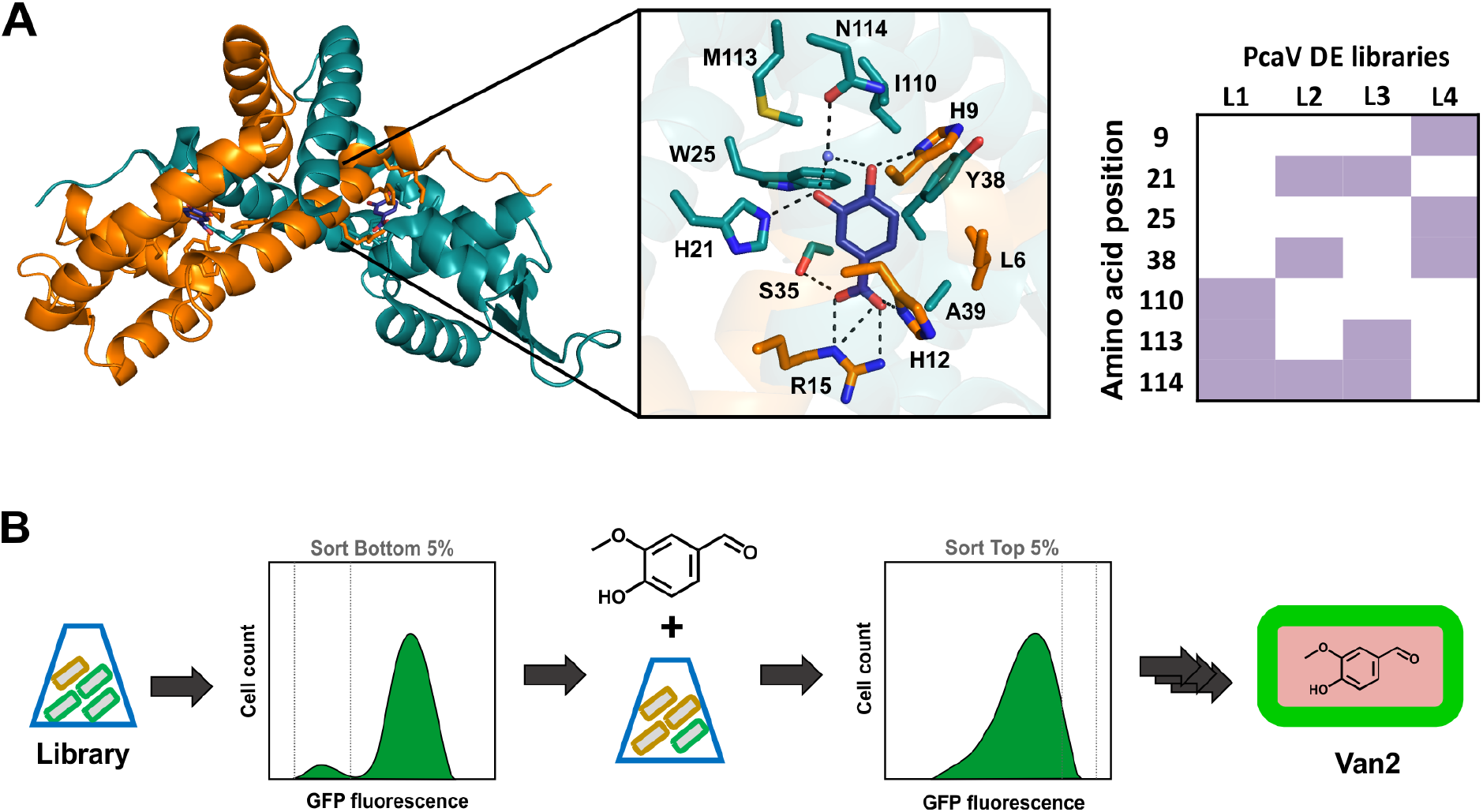
Directed Evolution of PcaV. **A.** The amino acids H9, H21, W25, Y38, I110, M113, and N114 in the PcaV effector binding site were selected for generation of completely randomized libraries, based on the PcaV-PCA structure (PDB: 4G9Y, 4FHT) [38]. **B.** Cultures of *E. coli* with the PcaV DE Libraries were used in a Fluorescence Activated Cell Sorting (FACS) counter-selection method for the enrichment of mutants with altered substrate specificity. Rounds of FACS were made alternating between negative selection of uninduced libraries with sorting of events on the bottom of the GFP fluorescent histogram and positive selection of induced libraries with sorting of events on the top of the GFP fluorescent histogram. Plate-based screening assay was performed on selected colonies from each sorted library and assessed for induction with the effector. The fluorescence was measured and the ratio induced/uninduced was used to select for responsive mutants.

The PcaV DE libraries were assembled (p15 plasmid), and transformed into electrocompetent *E. coli* cells harbouring the reporter (p44P_PV_-eGFP plasmid). The strain populations containing libraries 1 to 4 were combined and screened. Screening for functional mutants was performed using fluorescence-activated cell sorting (FACS), followed by plate screening and full titration analysis (**Fig. 2B**). The screening was performed to search for mutants able to detect for the industrial important chemicals, including vanillin (Table S7). To explore the potential to identify mutants with similar substrate recognition but different biosensor performance PCA was also included as a control.

The majority of variants in the mutant library were non-functional (unable to bind to the DNA) leading to eGFP expression in absence of any effectors, as observed from the population distribution in the FACS histogram for the uninduced library (**Fig. S2B**). The large number of non-functional mutants observed can be related to the allosteric regulation of PcaV, where small changes could lead to profound effects upon conformation impairing the DBD interaction with operator sequence [58]. Four rounds of FACS counter-selection was performed with successive negative and positive selection. For negative selection the library, in the absence of candidate effectors, was sorted to select for the sub-population with lowest fluorescence. For positive selection the library, in the presence of candidate effectors, was sorted for the sub-population with highest fluorescence (**Methods**). Functional mutants were selected from the directed evolution streams of PCA and vanillin induced libraries.

### *In vivo* performance of alternative PCA biosensors

From the PCA directed evolution screening, two mutants were identified with distinct features compared to the PcaV wild type. The mutant mutPCA1 had three amino acids substitutions (I110A, M113L, N114S) while the mutant mutPCA2 had two amino acids substitutions (W25Y, Y38W) (**Fig. S3A**). When tested with increasing concentrations of PCA the mutPCA1 system showed a higher basal expression and reduced dynamic range compared the wild type PCA biosensor (**Fig. S3A**). However, when assessed against other closely related candidate substrates, an indication that its substrate specificity pattern broadened towards benzoic acid and 2-hydroxybenzoic acid was observed (**Fig. S3B**). Analysing the modified amino acids of mutPCA1, the substitution of three amino acids on the upper side of the binding-pocket, including the change of the polar amide residue asparagine to the smaller polar alcohol residue serine (N114S), led to increased basal expression and broader specificity. In contrast the mutPCA2 biosensor system displayed a similar performance to the parental PCA biosensor (**Fig. S3A**). Analysing the modified amino acids of mutPCA2, inversion of the parental amino acids at position 25 and 38 was well tolerated leading to small increase of the basal expression and no change in the specificity.

### *In vivo* performance of the Van2 biosensor

From the vanillin directed evolution screening, the mutant Van2 was identified to respond to vanillin, but not the parent inducer PCA (**Fig. 3A**). Van2 was found to have three amino acids substitutions of I110V, M113S and A114N from the PcaV WT.

**Figure 3:**
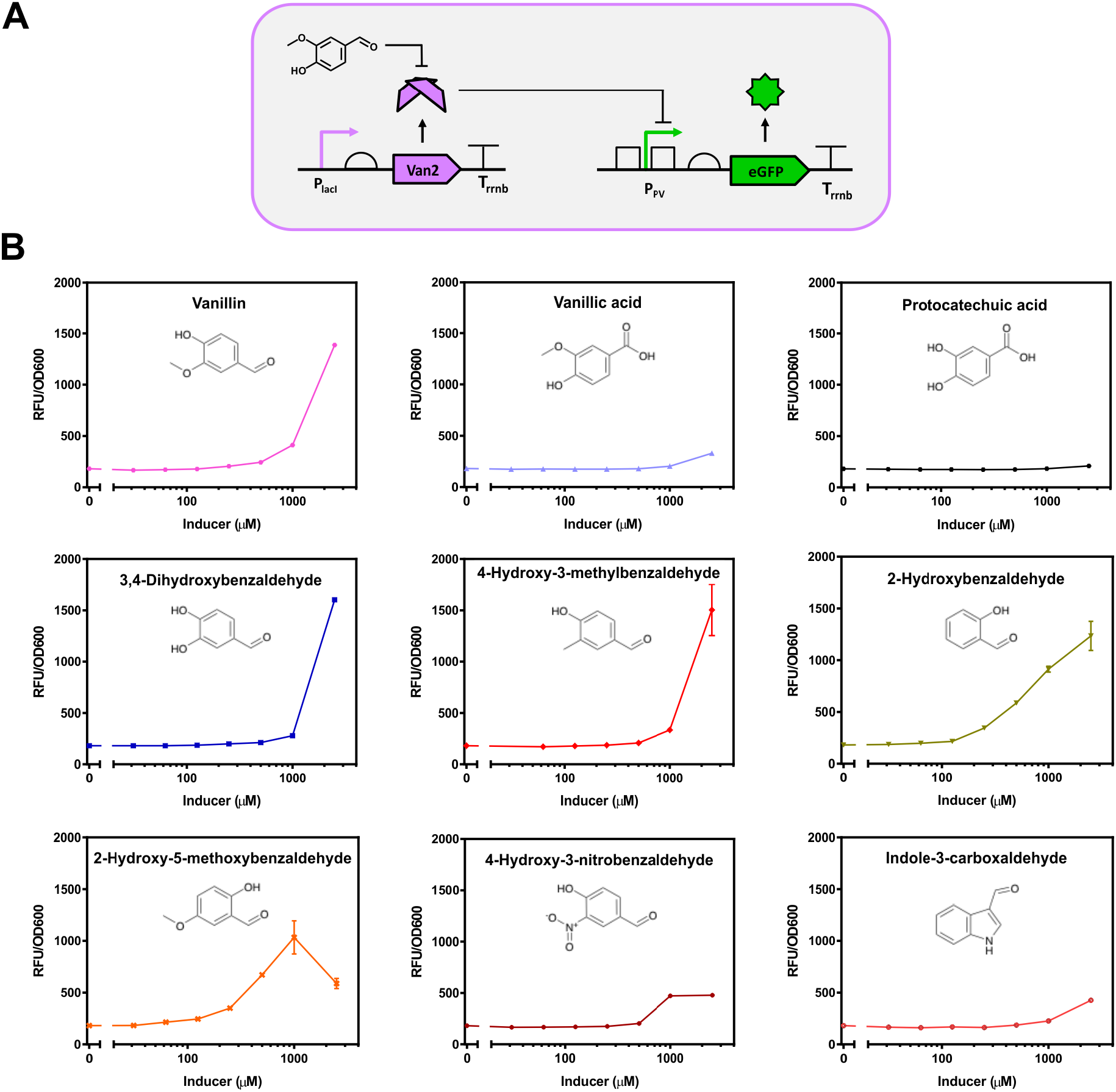
The Van2 Biosensor *in vivo* characterisation. **A.** Representation of the Van2 biosensor design. **B.** The Van2 Biosensor in *E. coli* RARE strain was induced with increasing concentrations of vanillin. The fluorescent gene expression normalised to cell density (RFU/OD_600_) was plotted. Each value represents the mean and standard deviation of 3 biological replicates.

Investigation of vanillin detection was made by GC-MS to determine if the signal was coming from vanillin itself or a metabolic conversion product by *E. coli*. The supernatant of cells exposed to vanillin revealed a substantial metabolism of vanillin to vanillyl alcohol (**Fig. S4A**). An induction test was then made with increasing concentrations of vanillyl alcohol and no fluorescent signal was observed for the latter, validating the vanillin detection by the Van2 system (**Fig. S4B**). Aldo-keto reductases and alcohol dehydrogenases have been reported to reduce aromatic aldehydes in *E. coli* [59, 60]. Indeed an engineered *E. coli* strain lacking all the genes for these enzymes, named RARE (reduced aromatic aldehyde reduction), was previously shown to reduce the conversion of benzaldehyde to benzyl alcohol [48]. Knockout strains of the individual genes coding for aldehyde reduction enzymes (*ΔdkgA, ΔdkgB, ΔyeaE, ΔyqhD, ΔyahK* and *ΔyjgB*) obtained from the Keio *E. coli* strain collection [61] and the RARE strain [48] were transformed with the Van2 biosensor system and tested with vanillin. All strains displayed enhanced output signal in response to vanillin compare to the wild-type, however the RARE strain had the best performance and was used for further performance characterisation of the Van2 biosensor (**Fig. S4C**).

Substrate specificity screen of the Van2 biosensor in the RARE strain was then assessed against 28 aromatic molecules, including other aldehydes, PCA and vanillic acid (**Table S8**). The Van2 biosensor detected vanillin with dynamic range of 7.7-fold and other related aromatic aldehydes (3,4-dihydroxyaldehyde, 4-hydroxy-3-methylbenzaldehyde and 2-hydroxybenzaldehide) with dynamic range higher than 6.0-fold at 2.5 mM (**Fig. 3B**).

### Van2 aTF biochemical and biophysical characterisation

In order to explore the altered substrate specificity observed *in vivo*, a series of biochemical and biophysical analyses were conducted to characterise the Van2 aTF. PcaV and Van2 proteins were expressed and purified (**Methods, Fig. S5A**). They were analysed to determine their behaviour in solution by size-exclusion chromatography coupled to multiangle light scattering (SEC-MALS) (**Fig. 4A**). Both PcaV and Van2 were observed as monodispersed (blue and red peaks respectively) with masses of 33.5 kDa for PcaV and 34.0 kDa for Van2 (blue and red strong lines inside the peaks) (**Fig. 4A**) indicating that they both form a homodimer in the absence of the effector, which is expected for functional aTFs [38].

**Figure 4:**
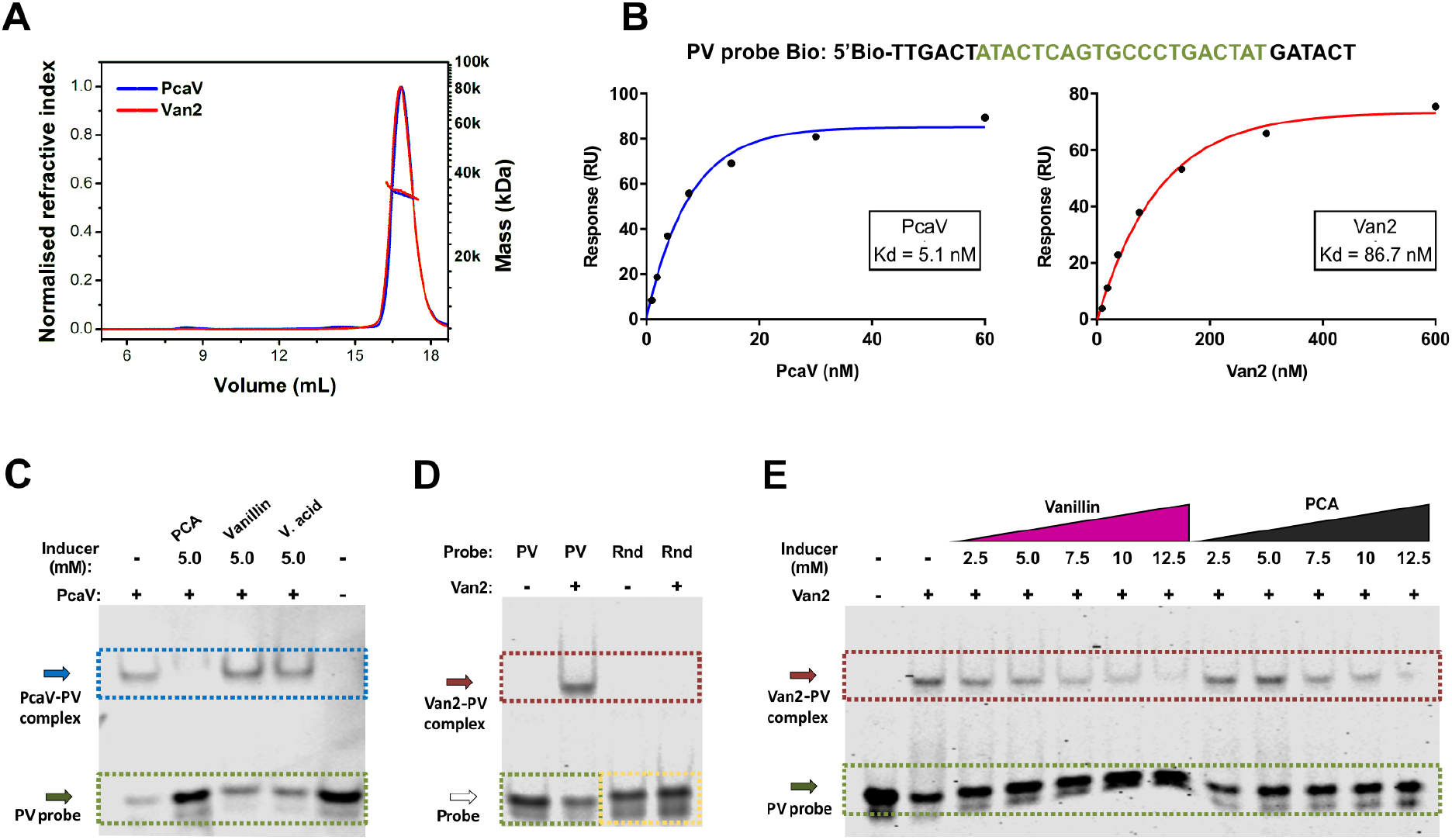
Biochemical and Biophysical characterisation of Van2. **A.** PcaV and Van2 analysis in solution by size-exclusion chromatography coupled to multi-angle light scattering (SEC-MALS). PcaV and Van2 traces are shown as monodispersed (blue and red peaks respectively) and the molecular weight at 34.0 kDa and 33.5 kDa (blue and red strong lines inside the peak) indicating formation of a homodimer. **B.** Surface Plasmon Resonance (SPR) of PcaV and Van2 interaction with the biotinylated DNA palindromic probe PV Bio. Concentrations of PcaV from 0.94 nM to 60 nM and Van2 ranging 9.4 nM to 600 nM were tested. The signal was plotted and a curve was traced to calculate the affinity constants (Kd). PcaV showed experimental affinity to the PV Bio probe with Kd of 5.05 nM. Van2 showed a Kd of 86.7 nM. **C to E** Electrophoretic Mobility Shift Assays (EMSA) of PcaV and Van2 with the 5’-infrared probe PV Bio. **C.** PcaV - PV probe complex is disrupted in the presence of 5 mM of PCA, but not with vanillin or vanillic acid at the same concentrations. **D.** Van2 forms complex with PV, but not with the Rnd probe containing the same nucleotides completely randomized. **E.** EMSA titrations with increasing concentrations of vanillin and PCA showed similar pattern of Van2-PV complex disruption.

The affinity of PcaV and Van2 aTFs to the palindromic DNA operator region was determined by Surface Plasmon Resonance (SPR) analysis using the DNA PV biotinylated probe (**Methods, Fig. 4B**). Concentrations of PcaV ranging from 0.94 nM to 60 nM and Van2 9.4 nM to 600 nM were tested to generate sensorgrams (**Fig. S5B**). The signal was plotted and a curve was traced to calculate the affinity constants (Kd). PcaV showed experimental affinity to the probe with a Kd of 5.1 nM, which is very close to the previously reported affinity for the Oi palindromic region in *S. coelicolor* (4.6 nM) [38]. Van2 showed a Kd of 86.7 nM, lower than the observed PcaV affinity (**Fig. 4B**). The lower affinity observed for Van2 compared to PcaV is also reflective of the *in vivo* performance of the Van2 biosensor, which shows a slightly higher basal expression (**Table 1**).

**Table 1.** Point mutational analysis of the PcaV to Van2 mutant.

| aTF | Amino acid position | Uninduced GFP/OD | Max/Min (2.5 mM induction) |  |  |
| --- | --- | --- | --- | --- | --- |
|  |  |  | Vanillin | PCA | Vanillic acid |
| **PcaV** | I110; M113; N114 | 34.2 ± 19.3 | 1.6 | 189.9 | 2.7 |
| **PcaV I110V** | **V110**; M113; N114 | 24.7 ± 6.5 | 1.7 | 304.4 | 8.2 |
| **PcaV N114A** | I110; M113; **A114** | 54.8 ± 10 | 2.5 | 18 | 2.2 |
| **PcaV M113S** | I110; **S113**; N114 | 90.1 ± 21.1 | 3.3 | 5.1 | 4.9 |
| **PcaV I110V N114A** | **V110**; M113; **A114** | 74.8 ± 7.8 | 3.5 | 10.7 | 2.4 |
| **PcaV I110V M113S** | **V110**; **S113**; N114 | 561.8 ± 52.9 | 3.3 | 2.4 | 3.9 |
| **PcaV M113S N114A** | I110; **S113**; **A114** | 1830.1 ± 60.1 | 3.1 | 1.1 | 1.6 |
| **Van2** | **V110**; **S113**; **A114** | 253.4 ± 0.3 | 5.8 | 1.2 | 2.2 |

Electrophoretic Mobility Shift Assays (EMSA) of PcaV (**Fig. 4C**) and Van2 (**Fig. 4D and 4E**) were next performed with the 5’-infrared tagged DNA PV probe in order to determine the interaction of the aTFs with the DNA operator region in the absence and presence of the effectors. PcaV formed a complex with the PV probe and this complex was disrupted in the presence of 5 mM of PCA, but not with vanillin or vanillic acid at the same concentration (**Fig. 4C**). Van2 formed a complex with the PV probe, but not with the Rnd probe containing the same nucleotides as PV completely randomized (**Fig. 4D**). EMSA titrations with increasing concentrations of effectors, from 2.5 mM to 12.5 mM, showed Van2-PV complex disruption for vanillin, but also PCA (**Fig. 4E**). The EMSA results indicate that *in vitro* complete derepression of PcaV-PV occurs with PCA, in contrast no derepression is observed with vanillin which is consistent with the observed *in vivo* biosensor performance. Van2 responds to vanillin but is still responsive to PCA in contradiction with the observed *in vivo* biosensor performance. The Van2 effector response pattern has changed compared to the WT PcaV, where partial derepression of the Van2-PV complex occurs. This altered response pattern is reflective of the *in vivo* performance of the Van2 biosensor that shows a reduced maximal expression signal for vanillin and no expression signal for PCA.

### Point mutational analysis of the PcaV to Van2 mutant

In order to investigate the key amino acid substitutions responsible for the altered *in vivo* specificity, a strategy was employed to create and assess the point mutations of each amino acid in the mutant Van2. Intermediate mutants with one or two point mutations ranging from PcaV (I110, M113, N114) to Van2 (V110, S113, A114) were generated by site-directed mutagenesis. The mutants were tested in the biosensor system with increasing concentrations of vanillin, PCA and vanillic acid (**Fig. 5A**). The substitution of the amino acid I110 individually (PcaV I110V) did not affect the performance of the biosensor or the substrate specificity. The substitution of either amino acids at positions N114 or M113 individually (PcaV N114A or PcaV M113S), or of M113 in combination with I110 (PcaV I110V and N114A), drastically reduced the signal output, but maintained substrate specificity toward PCA. The substitution of I110 and M113 in combination (PcaV I110V M113S) showed a small increase in basal expression and was partially responsive to all the substrates assessed (vanillin, vanillic acid and PCA). The substitution of M113 and N114 in combination (PcaV M113S N114A) displayed a high basal expression indicating reduction of DNA binding affinity, but a complete change of specificity. Similar to Van2 it was responsive to vanillin, partially responsive to vanillic acid and unresponsive to PCA. The additional substitution I110V that leads to Van2 (I110V M113S N114A), conferred reduction of the basal expression whilst maintaining the substrate specificity pattern. Interestingly, this same amino acid substitution in PcaV I110V had no significant effect on the performance compared to PcaV (**Fig. 5A, Table 1**).

**Figure 5:**
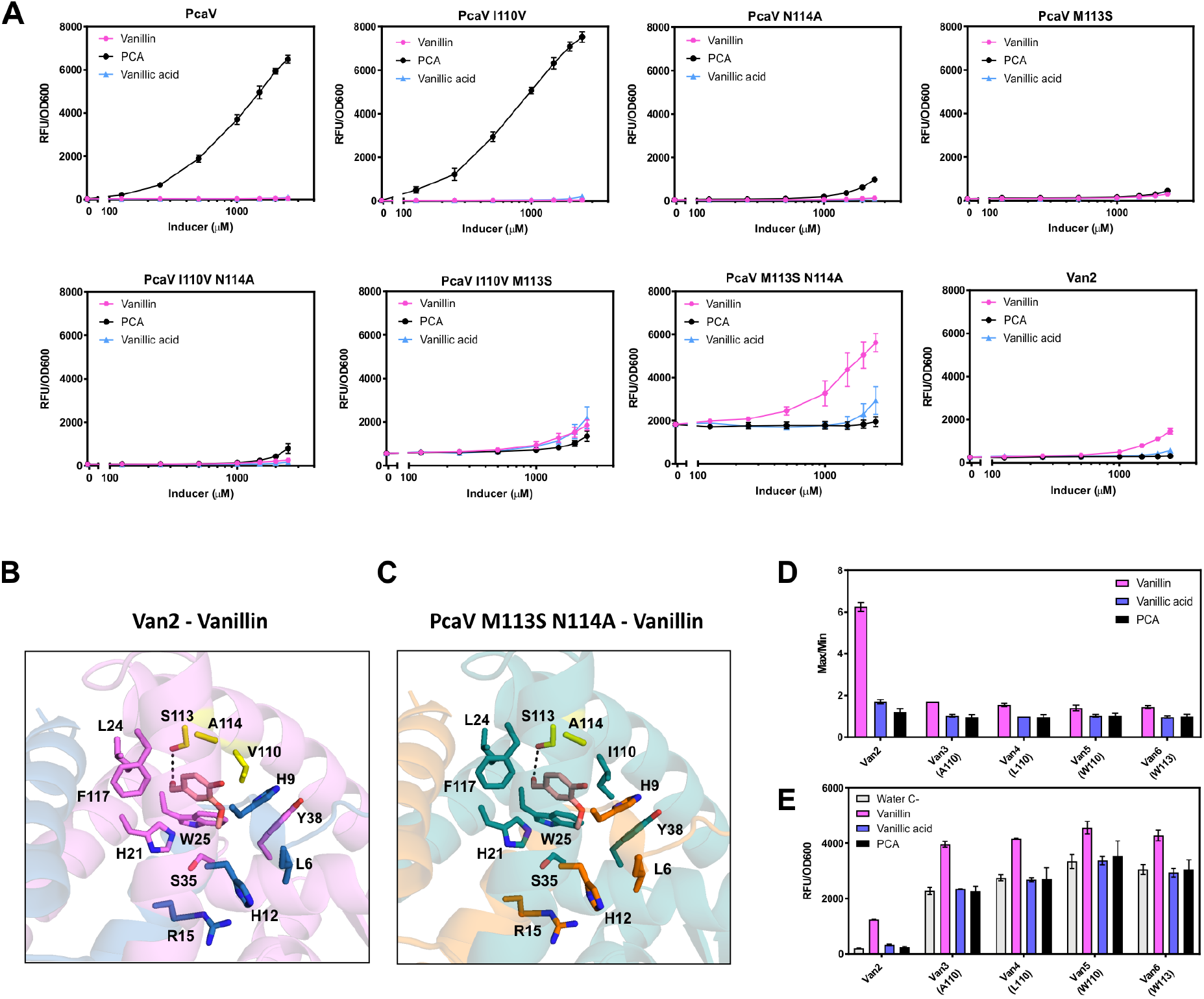
Test of mutants from PcaV to Van2 and the Van mutants A. Point mutants of the 3 amino acid positions ranging from PcaV (I110, M113 and N114) to Van2 (V110, S113, A114) were generated and all systems were induced with increasing concentrations of vanillin, PCA and vanillic acid. PcaV I110V kept the PCA detection performance and substrate specificity pattern. PcaV N114A, PcaV M113S and PcaV I110V N114A led to reduced PCA detection without substrate specificity change. PcaV I110V M113S led to reduced PCA detection and substrate specificity change with detection of vanillin and vanillic acid. PcaV M113S N114A showed complete substrate specificity change being responsive to vanillin, partially responsive to vanillic acid, and unresponsive to PCA, however the system had a very high basal expression. B. Homology modelling and docking of Van2 and the double mutant PcaV M113S N114A. Docking of the models for Van2 and PcaV M113S N114A with vanillin showed, in the 2^nd^ best docked conformation, an interacting with S113 of Van2 via a hydrogen bond between the aldehyde functional group of vanillin and the hydroxyl group of the serine. This conformation was only observed for Van2 and PcaV M113S N114A, mutants that displayed vanillin detection *in vivo*. C. Further mutations of Van2 to explore further amino acids substitutions at positions 110 and 113 were induced with vanillin, PCA and vanillic acid at 2.5 mM. Presence of apolar small, medium and bulky amino acids at 110 on Van3, Van4 and Van5 respectively, as well as a bulky amino acid at 113 on Van6 kept a high basal expression of the systems leading to low max/min. The fluorescent gene expression normalised to cell density (RFU/OD_600_) is shown. Each value represents the mean and standard deviation of 3 biological triplicates.

### PcaV to Van2 modelling and docking

Homology modelling and docking of the generated mutants were made to help understand the role of amino acid substitutions upon the altered aTF specificity. Docking of Van2 with vanillin (**Methods**) led to the lowest energy conformation (Score 1: ΔG_binding_ = −6.4 kcal/mol) that had the same location as PCA in the PDB crystal structure (4FHT) (**Table S9**). However interestingly, the next lowest conformation (Score 2: ΔG_binding_ = −6.1 kcal/mol) had vanillin in a different position interacting with S113 of Van2 via a hydrogen bond between the aldehyde functional group of vanillin and the hydroxyl group of the serine (**Fig. 5B**). A similar effector molecule orientation was also observed for the docking of PcaV double mutant M113S N114A with vanillin (**Fig. 5C**), consistent with the observed *in vivo* response (**Fig. 5A**). As a control re-docking of PcaV with PCA was performed, this indicated the lowest energy conformation state (Score 1: ΔG_binding_ = −7.6 Kcal/mol) with PCA positioned in the same orientation as described in the crystal structure for PcaV-PCA complex (4FHT) (**Fig. S6**). These docking conformations are consistent with the observed *in vivo* biosensor response. Specifically the amino acids S113 and A114 appear to play a role in the detection of vanillin by the PcaV double mutant (M113S N114A) and by Van2.

### Further point mutation of Van2 biosensor

Further mutants of Van2 were designed in an attempt to more extensively explore the role and dependency of the amino acid position at 110. The strategy was made based on the *in vivo* observation that S113 and A114 are essential for specificity change, but the V110 alone did not affect the aTF interaction with PCA. Three mutations were incorporated into Van2 at position 110, changing for nonpolar amino acids with increasing side chain size, generating Van3 (A110), Van4 (L110) and Van5 (W110). Additionally the Ser113 position was swapped to a nonpolar bulky residue, to generate Van6 (W113) (**Table 2**). The Van3-6 biosensors all displayed a high basal expression in the absence of effector, indicative of impaired DNA binding. However, this basal expression appears to be lower for the mutants with smaller amino acids (Van3 and Van 4) and higher for those with bulky amino acids (Van5 and Van6) (**Table 2**). The overall performance for vanillin detection was lower for all the new Van aTF mutants (<2-fold) compared to Van2 (>6-fold) (**Fig. 5D, Table 2**). This result indicates that Val110 plays an important role in the stabilization of the Van2-DNA interaction, whilst still allowing detection of vanillin. The mechanistic basis of this interaction is not intuitive though, as the presence of other apolar amino acids such as I110 in PcaV, A110 in Van3 and L110 in Van4 led to high basal expression. Overall, these results indicate that the repression/derepression mechanism behind the Van2 induction by vanillin depend on the presence of the all three mutated amino acids.

**Table 2.** Further point mutation of Van2 biosensor:

| aTF | Amino acid position | Mutated amino acid | Uninduced GFP/OD | Max/Min (2.5 mM induction) |  |  |
| --- | --- | --- | --- | --- | --- | --- |
|  |  |  |  | Vanillin | PCA | Vanillic acid |
| **Van2** | V110; S113; A114 | X | 199.4 ± 5.9 | 6.25 | 1.20 | 1.70 |
| **Van3** | **A110**; S113; A114 | Nonpolar small side chain | 2280.6 ± 131.2 | 1.70 | 0.97 | 1.03 |
| **Van4** | **L110**; S113; A114 | Nonpolar medium side chain | 2748.7 ± 114.0 | 1.55 | 0.97 | 1.00 |
| **Van5** | **W110**; S113; A114 | Nonpolar bulky side chain | 3339.0 ± 249.3 | 1.40 | 1.03 | 1.03 |
| **Van6** | V110; **W113**; A114 | Nonpolar bulky side chain | 3047.5 ± 175.9 | 1.45 | 1.00 | 0.97 |

## Discussion

The development of new aTF-based biosensors is an important area of the synthetic biology field, as aTFs are versatile biological tools for a range of methodologies and applications [23, 24]. Developing biosensors for aromatic molecules is especially important due to their utility in industry and their availability from biomass sources [62]. Use of natural sources of sensing machinery to generate biosensors has been an approach successfully applied with a range of examples in literature [1, 14], however it relies on a limited set of aTFs with known inducer [7]. Therefore strategies to develop *de novo* aTFs for desired effectors are essential. In this sense, the directed evolution of aTFs to change the specificity patterns is an interesting strategy, regardless of the challenges associated to the allosteric nature of these proteins.

The PCA biosensor in *E. coli* developed here employed a heterologous aTF from *Streptomyces coelicolor* along with engineered chimeric promoter-operators based on phage promoter architecture. This approach yielded the highly responsive P_PV_ version of the PCA biosensor for detection of meta, para or di-hydroxyl substituted benzoic acid. The observed large dynamic range (>150-fold) is useful for selection methods based on the separation of sub-populations from genetic libraries, for example by FACS. These characteristics in conjunction with the available crystal structure of PcaV paved the directed evolution strategy to change the substrate specificity pattern of PcaV.

To our knowledge, this is the first work on the directed evolution of a MarR family allosteric transcription factor. In contrast to other aTF families, the MarR family has a particular structure with the DBD and EBD fused to each other [1, 41], and amino acids from different monomers are contained in the EBD [42]. A conserved scaffold is shared by this family, however the repression mechanism is variable among its members [42]. The PcaV mechanism was proposed based on the observation of apo and PcaV-PCA complex structures, where the PCA binding induces rotation of the DBD leading to a conformation incompatible with DNA binding [38]. However, there are limited examples of MarR-DNA complex structures available [41], which is also true for PcaV [38]. Taking the allosteric nature of aTFs, this limited structural knowledge makes it even harder to predict the effect of amino acid substitutions upon the DNA binding affinity of the aTF [58]. As observed here by examining the broad fluorescence distribution of the uninduced library, it can be seen that DNA binding was affected although only amino acid positions in the EBD were targeted (**Fig. S2B**). The inherent allosteric mechanism of aTFs can hamper the search of functional mutants with the desired new property from genetic libraries [58, 63]. High throughput methods coupled with rounds of negative (for functional DNA binding) and positive (for binding of the new effector) counter-selection have previously been employed to address these challenges by removing non-functional mutants and providing enrichment of desired mutants in the directed evolution of AraC [26, 36], PobR [28] and TrpR [29] for example. Other studies have combined high throughput screening and selective pressure methods in order to remove non-functional mutants [27, 64]. In this study, the PcaV directed evolution was performed using a FACS counter-selection method with four rounds of sorting to permit identification of the Van2 aTF.

The Van2 biosensor detected vanillin and other aromatic aldehydes *in vivo* without responding to the parental inducer PCA. The vanillin metabolism by the biosensor host strain was analysed by GC-MS, which showed vanillin was partially reduced to vanillyl alcohol, but subsequent assessment of the alcohol against the biosensor indicated no detection. Furthermore, the Van2 system in the *E. coli* RARE strain showed the greatest signal output and dynamic range in response to vanillin, indicative of a higher availability of the effector inside the cell. *In vitro* a small reduction of the binding affinity of Van2 aTF to the PV DNA operator (86.7 nM) was observed compared to PcaV (5.05 nM) assessed by SPR, which was consistent with the higher basal expression shown *in vivo* for the Van2 biosensor. As observed on the EMSA assays, Van2 responds to vanillin and was still responsive to PCA *in vitro*. However, PcaV was not responsive to vanillin and was very responsive to PCA. The absence of an *in vivo* response to PCA by the Van2 biosensor could most likely be explained by physiological processes such as membrane transport selectivity affecting the intracellular availability of the effector inside the cell.

The amino acid mutations of PcaV to Van2 (I110V, M113S and N114A) led to a change in the substrate specificity. Individual mutations from the parent PcaV to Van2 indicated that the amino acids S113 and A114 have a major role on the detection of vanillin and the amino acid V110 is important to maintain repression in the absence of effector, most likely indicating that V110 is required for functional DNA binding. Interestingly, homology modelling and docking of the active *in vivo* mutants, Van2 and PcaV M113S N114A, with vanillin, indicated an interaction between the aldehyde functional group of vanillin and the serine at position 113. A similar observation was reported in the directed evolution of NitR to detect ε-caprolactam [65]. Homology modelling and docking of the NitR-L117F mutant showed a shorter distance between the effector and the mutated amino acid inside the binding-pocket compared to the WT, which suggested an association with the improved sensing [65]. Therefore the docking results for Van2 could indicate an association between the altered specificity pattern in Van2 with the availability of the position of the new effector in the binding pocket.

Other genetically encoded biosensors for aromatic aldehydes have been reported [66–71]. Among transcriptional based biosensors, a system based on the BldR aTF of archea *Sulfolobus solfataricus* was developed in *E. coli* for the detection of benzaldehyde [66]. Another study used a computational based approach to reengineer the qacR aTF and generate a vanillin biosensor [67]. In other studies, biosensors based on the EmrR aTF from *E. coli* were developed allowing detection of vanillin, however these biosensors were also shown to be responsive to other aromatic acids, including vanillic and coumaric acid [68–70]. A more recent study generated a biosensor in *E. coli* based on the native YqhC aTF for the detection of aliphatic and aromatic aldehydes including glycoaldehyde and vanillin [71]. The Van2 biosensor developed here displayed high dynamic range with up to 7.7-fold signal, showed specificity for detection of vanillin and other aromatic aldehydes, and was unresponsive *in vivo* to the parental aromatic acid effector, PCA.

## Conclusion

We report a new aTF-based biosensor to detect aromatic aldehydes, based on the reengineering of the PcaV aTF. Van2 was characterised and the role of the amino acid substitutions that led to vanillin sensing were assessed *in vivo*. The Van2 biosensor can detect vanillin and 3,4-dihydroxybenzaldehyde used in the flavors industry and as chemical building blocks respectively [47, 72]. It could be applied for the improvement of biotechnological production of high value chemicals [48] and enable screening methodologies for discovery of enzymes able to bioprocess renewable lignin sources [68].

This study expands the knowledge upon the directed evolution for biosensors, specifically exploring the plasticity of a member of the MarR family of transcription factors. We demonstrate the directed evolution of the PcaV aTF to change its substrate specificity pattern and generate a biosensor for aromatic aldehydes.

## List of abbreviations

aTF: allosteric transcription factor
PCA: protocatechuic acid
EBD: effector binding domain
DBD: DNA binding domain
RNAP: RNA polymerase
DMSO: dimethyl sulfoxide
RFU: relative fluorescence unit
OE-PCR: overlap-extension polymerase chain reaction
DE: directed evolution
FACS: fluorescence activated cell sorting
MTBE: methyl-tert-butyl ether
EMSA: electrophoretic mobility shift assay
SEC-MALS: size-exclusion chromatography coupled to multi-angle light scattering
SPR: Surface Plasmon Resonance

## Declarations

### Acknowledgments

We would like to thank Dr. Joanne Porter from Prof. Nick Turner’s group for the support with the GC-MS analysis; Prof. David Leys’ group for the help on the protein purification and Dr. Thomas Jowitt from the Biomolecular Analysis Core Facility of the Faculty of Biology, Medicine and Health for the SEC-MALS and SPR analysis. We also would like to thank Dr. Linus Johannissen and Dr. Sam Hay for the valuable advice and discussion regarding the modelling and docking methods.

### Funding

ND is supported by the BBSRC David Phillips Fellowship (BB/K014773/1) and BBSRC grant (BB/P01738X/1). LM is supported via Science without Borders/Ciências sem fronteiras scheme from the CNPq, Brazil (233608/2014-1).

### Availability of data and materials

The datasets during and/or analysed during the current study available from the corresponding author on reasonable request.

### Authors’ contributions

ND designed and coordinated the study. LM planned and performed the experiments. AC supported the directed evolution design. LM and ND analysed the data. LM and ND wrote the manuscript. All authors read and approved the final version of the manuscript.

### Competing interests

The authors disclose no conflicts.

